# You shall know a species by the company it keeps: leveraging co-occurrence data to improve ecological prediction

**DOI:** 10.1101/2023.02.15.528518

**Authors:** Andrew Siefert, Daniel C. Laughlin, Francesco Maria Sabatini

## Abstract

1. Making predictions about species, including how they respond to environmental change, is a central challenge for ecologists. Due to the huge number of species, ecologists seek generalizations based on species’ traits and phylogenetic relationships, but the predictive power of trait-based and phylogenetic models is often low. Species co-occurrence patterns may contain additional information about species’ ecological attributes not captured by traits or phylogenies.
2. We propose using ordination to encode the information contained in species co-occurrence data in low-dimensional vectors that can be used to represent species in ecological prediction. We present an efficient method to derive species vectors from co-occurrence data using GloVe, an unsupervised learning algorithm originally designed for language modeling. To demonstrate the method, we used GloVe to generate vectors for nearly 40,000 plant species using co-occurrence statistics derived from a global vegetation dataset and tested their ability to predict elevational range shifts in European montane plant species.
3. Co-occurrence-based species vectors were weakly correlated with traits or phylogeny, indicating that they encode unique information about species. Models including co-occurrence-based vectors explained twice as much variation in species range shifts as models including only traits or phylogenetic information.
4. Given the widespread availability of species occurrence data, species vectors learned from co-occurrence patterns are a widely applicable and powerful tool for encoding ecological information about species, with many potential applications for describing and predicting the ecology of species, communities, and ecosystems.

## INTRODUCTION

Ecologists face the daunting challenge of making predictions about species, such as where they occur, how they perform, how they respond to environmental change, how they impact ecosystem functioning, and how they interact with other organisms (Guisan & Zimmermann, 2000; Strydom et al., 2021). This task is made difficult by the sheer diversity of life on Earth. Despite the huge amount of biodiversity data currently available, we know little to nothing about the ecology of most species (Hortal et al., 2015). Finding ways to generalize—to take information learned from the species that have been studied and apply it to other species and contexts—is therefore a necessity.

Using species’ traits is one approach to meet the challenge of prediction in ecology (Lavorel & Garnier, 2002; McGill et al., 2006). The benefits of a trait-based approach are that it allows us to make predictions that are mechanistic, because traits capture how species function, and general, because we assume that species with similar trait values behave similarly. There has been tremendous progress over the past 20 years in collecting trait data and assembling global trait data sets (Jones et al., 2009; Kattge et al., 2020; Madin et al., 2016), so the availability of trait data is increasing, but large gaps remain. There have been many efforts to use traits to predict species’ distributions, abundances, demographic performance, responses to environmental change, and effects on ecosystem functioning (Angert et al., 2011; Funk et al., 2017), but the amount of variation explained by traits is often low (Beissinger & Riddell, 2021; Paine et al., 2015), suggesting that measured traits often do not capture all relevant aspects of species’ ecology. Using phylogenetic relatedness as a proxy for unmeasured traits is one way to get around the issue (Cadotte et al., 2013; Webb et al., 2002), but this approach assumes that those traits are phylogenetically conserved. So, while trait-based and phylogenetic approaches are promising, they have clear limitations. To make better predictions, it is necessary to capture the ecological characteristics of species that traits and phylogenies miss.

Species co-occurrence patterns reflect many ecological characteristics of the species involved (Fig. 1a-b), including their environmental tolerances, resource requirements, competitive effects and responses, susceptibility to enemies, associations with mutualists, and dispersal ability (Elton, 1946; Peres-Neto et al., 2001). Many of these characteristics may not be captured by easily measured traits or phylogenetic relationships. Ecologists have long recognized that species co-occurrence patterns contain valuable ecological information, and there is a long history of attempts to use them to infer ecological processes (Diamond, 1975; Gotelli, 2000), generating an equally long-running debate over the validity of these attempts (Connor & Simberloff, 1979; Lewin, 1983). We argue that the very property of species co-occurrence patterns that challenges their usefulness for inference—i.e., they contain the signals of multiple ecological processes—is what makes them a potentially powerful tool for prediction. Encoding the information in species co-occurrence patterns and including it in models may improve our ability to make predictions about species (Carmona & Pärtel, 2021).

**Figure 1.**
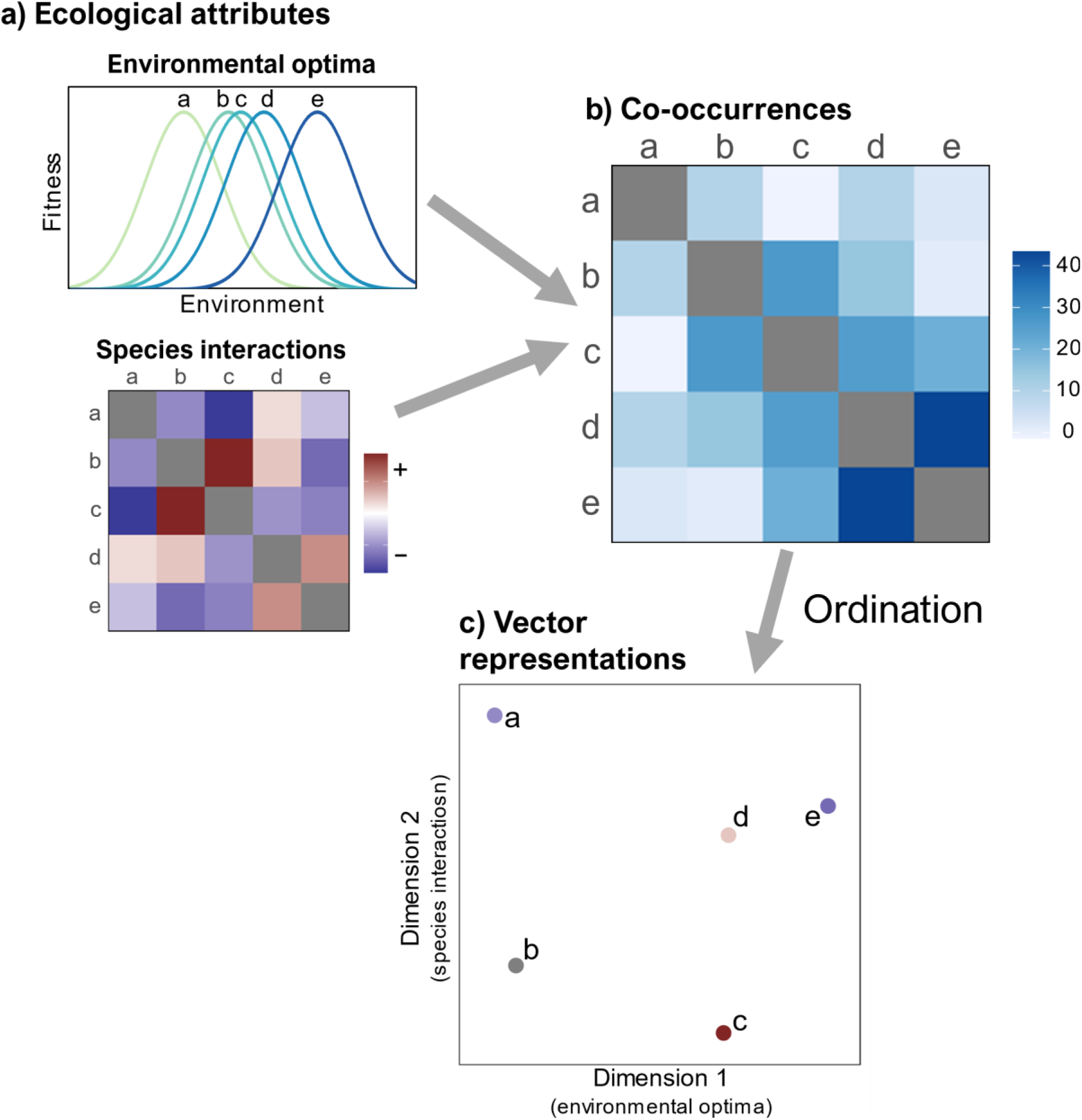
Conceptual illustration of how information about species’ ecological attributes contained in co-occurrence patterns are encoded in vector representations of species. a) In this hypothetical example using simulated data, five species are characterized by their responses to an environmental gradient and pairwise interaction strengths, ranging from facilitation (positive) to competition (negative). b) These attributes drive species co-occurrence patterns, such that species with similar environmental optima and positive interactions (e.g., species d and e) are more likely to co-occur frequently. c) Ordination of the co-occurrence data results in vector representations of species, such that species that co-occur more frequently are close to each other in the vector space. These vectors encode information about the ecological processes that produced the co-occurrence patterns. Here, dimension 1 primarily reflects species’ environmental optima and dimension 2 primarily reflects species interactions. Points representing species are colored according to their interactions with species b to highlight that species with positive interactions (e.g., species b and c) are close to each other along Dimension 2, and species that have negative interactions (e.g. species a and b) are far apart.

To capture this information, we propose borrowing an idea from language models. Just as species co-occurrences contain information about species’ ecology, word co-occurrences contain information about word meaning, a principle that Firth (1957) famously summarized as “you shall know a word by the company it keeps”. Language models seek to capture this information in word embeddings—representations of words as numerical vectors that encode information about their meaning (Hinton, 1986; Li & Yang, 2018). These vectors can be learned by training a model to predict word co-occurrences (e.g., the frequency with which a word appears in the context of a target word) in a body of text (Collobert et al., 2011). The output of interest from the model is not the predicted co-occurrence frequencies themselves, but rather the internal representations of words (i.e., vectors) that the model learns in the course of generating the predictions. Information about word meaning contained in the co-occurrence patterns is encoded in the vectors, such that words with similar meaning will have vectors with similar numerical values. Word embedding models are most successful when they are trained on huge datasets with billions of words (Mikolov et al., 2013). The resulting vectors can be used to represent words in a variety of applications, such as document classification, sentiment analysis, text summarization, and translation. This is an example of transfer learning—taking information learned from one context (i.e., predicting word co-occurrence frequencies) and applying it to others (e.g., document classification; Pan & Yang, 2010).

We propose that a similar approach can be applied to species. The widespread availability of species occurrence data means that it is possible to calculate co-occurrence statistics for many species, and the ecological information contained in these statistics can be encoded in vectors to represent species in ecological prediction (Fig 1c). Ecologists have long used ordination techniques to encode the information in high-dimensional community data in low-dimensional vectors (e.g., species scores or loadings). Several authors have recently proposed adapting word embedding algorithms to generate “species embeddings” that capture information about species’ ecology (Angelov, 2018; Chen et al., 2017; Joseph, 2020), but these approaches have not been fully leveraged for ecological prediction. To do so, we propose deriving vector representations of species from large, global co-occurrence datasets and using these vectors in place of or in addition to traits and phylogenies to represent species in ecological prediction tasks.

Here, we describe an efficient approach for learning vector representations of species from co-occurrence data using GloVe (Pennington et al., 2014), an unsupervised learning algorithm originally created to generate word embeddings. Second, we present a case study in which we use GloVe to generate vectors for nearly 40,000 plant species using a global vegetation database (Sabatini et al., 2021). We test the ability of these vectors to predict species’ range shifts using an independent data set of plant species in European mountains (Lenoir et al., 2008) and compare the predictive performance of co-occurrence-based vectors derived using GloVe and a traditional ecological ordination method (transformation-based PCA) with that of functional traits and phylogenetic information. Finally, we discuss potential extensions and applications of this approach in ecology.

## METHODS

### The framework

The general use case for our approach is a situation in which an ecological attribute of interest (“prediction target”, e.g., fitness, range shift, environmental response or effect) has been measured on a subset of species (“observed species”) and the goal is to predict this attribute for other species (“unobserved species”) for which the prediction target has not been directly measured. Our general framework is to 1) obtain a species occurrence dataset that includes both the observed and unobserved species and calculate species co-occurrence statistics, 2) apply an ordination method to the co-occurrence data to derive low-dimensional vector representations of the species, 3) fit a model of the prediction target using the vector values of the observed species as predictors, and 4) use this model to make predictions for the unobserved species based on their vector values. Ideally, a very large occurrence dataset would be used to derive the species vectors to maximize their ecological information content. In theory, any ordination method could be used to derive the species vectors, but in practice most ordination methods traditionally used in ecology may not be able to handle very large species occurrence datasets, creating a need for efficient, scalable methods.

### GloVe algorithm

The GloVe (Global Vectors for Word Representation) algorithm (Pennington et al., 2014) is a method for learning word embeddings from word co-occurrence statistics. The model is trained on a word-word co-occurrence matrix whose entries *X_ij_* count the number of times word *i* appears in the context (i.e., within a certain number of words) of word *j* within a body of text. When applied to species, *X_ij_* is the number of times species *i* co-occurs (e.g., is found in the same plot) with species *j*. The objective of the algorithm is to learn vectors *w_i_* for the “target” species and 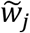 for the “context” species such that their dot product equals the logarithm of the species’ cooccurrence frequency:

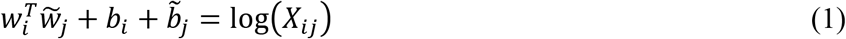

where *b_i_* and 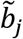 are species-specific intercepts (also known as “bias” terms). The algorithm learns two vectors for each species, *w* and 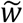. When the co-occurrence matrix is symmetric, the two vectors are equivalent and only differ due to their random initializations when fitting the model. Equation 1 may be seen as a matrix factorization or dimensionality reduction problem, in which an *n*-species by *n*-species co-occurrence matrix *X* is reduced to an *n*-species by *m*-dimension matrix *W*, where *m* is the length of the species vectors as defined by the user.

A drawback of the matrix factorization approach is that it gives equal weight to all co-occurrences, even those that co-occur rarely, which may be noisy and carry less information than the more frequent co-occurrences. Pennington et al. (2014) proposed including a weighting function and casting equation 1 as a weighted least squares regression in which the training objective is to find the weights (*w* and 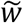) and biases (*b* and 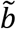) that minimize the following loss function:

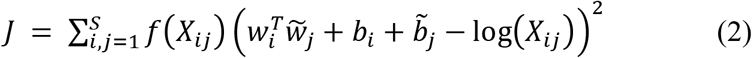

where *S* is the number of species and *f*(*X_ij_*) is a weighting function that puts less weight on rare co-occurrences. Pennington *et al*. (2014) proposed the weighting function:

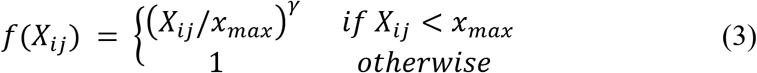

where *X_max_* is a threshold of co-occurrence counts below which co-occurrences are given less weight, and *γ* is a parameter that controls how quickly the weight decreases for co-occurrence counts below the threshold.

A key feature of the GloVe algorithm is that it is only trained on non-zero elements of the co-occurrence matrix. The reasoning is that non-co-occurrences (i.e., word pairs that never co-occur) carry less information than co-occurrences, and because word co-occurrence matrices are typically sparse (mostly zeroes), training on the non-zero elements reduces training time and makes the algorithm more efficient. We argue that the same reasoning applies to species co-occurrences, so we propose training the algorithm on non-zero entries of a species co-occurrence matrix. It may be possible that non-co-occurrences contain valuable information about species and that including this information could improve the performance of the model. This is something worth exploring in the future, though we note that the GloVe algorithm would need to be modified since the logarithm of 0 is undefined.

To generate word embeddings, GloVe is typically trained on word co-occurrence counts derived from a large body of text such as Wikipedia or collections of millions of news articles. The model may also be trained on a domain-specific corpus with the aim of creating word vectors that encode information specifically relevant to that domain. The vector dimension *m* and weighting function parameters *X_max_* and *α* are not estimated from the data but rather must be chosen. The word vectors themselves are estimated using an optimization routine, such as stochastic gradient descent, where the objective is to minimize the cost function in equation 2. The real test of model performance is not the accuracy of the predictions of co-occurrences, but rather the performance of the word vectors in external prediction tasks. Similarly, when generating species vectors, we are primarily interested in their ability to make ecological predictions about species.

### Case study

Here we present a case study demonstrating how to generate vector representations of species from co-occurrence data and how the vectors can be used in an ecological prediction task. We focus on predicting species range shifts in response to climate change, as this is one of the most urgent prediction tasks facing ecologists. Briefly, we generate species vectors of varying dimension using two ordination methods, GloVe and transformation-based principal component analysis (tb-PCA), on a global vegetation plot database (Sabatini et al., 2021) and test the ability of the vectors to predict elevational range shifts of 162 plant species in European mountains using the dataset of Lenoir *et al*. (2008; “range shifts dataset” hereafter). We compare the predictive performance of models using co-occurrence-based species vectors, functional traits, and phylogenetic relationships.

#### Generating species vectors

We derived species vectors using species co-occurrence counts calculated from sPlotOpen (Sabatini et al., 2021), an environmentally balanced global dataset of vegetation plots. sPlotOpen consists of three resampled subsets of the full sPlot database (Bruelheide et al., 2019). We combined all three subsets, resulting in a plot selection that may be less environmentally balanced due to overlap across the resamples. We removed taxa that did not have full binomial names or only occurred in one plot, resulting in a dataset containing 93,086 plots, 39,905 species, and 1,839,072 plot-level species occurrences. In addition to this global occurrence dataset, we created a “local” occurrence dataset that only included plots containing at least one species present in the range shifts dataset. The rationale for this is that we wanted to compare the predictive performance of vectors trained on global co-occurrences with those trained on “local” co-occurrences only, which contain less total information but potentially a higher proportion of information relevant to the prediction task. The “local” co-occurrence dataset contained 26,011 plots, 15,788 species, and 699,573 plot-level species occurrences. We created species co-occurrence matrices, whose entries were the number of plots in which a given species pair co-occurred, using each dataset.

We generated species vectors by training the GloVe algorithm on the non-zero entries of the species co-occurrence matrices. We fit models using different combinations of vector dimension (*m* = 4, 8, 16, 32, or 64) and weighting function cutoff (*x_max_* = 1 or 100). We set γ = 0.75 for all model runs, following Pennington *et al*. (2014). We trained the models using Adam, a stochastic gradient descent optimizer, with an initial learning rate of 0.001, for up to 50 epochs (iterations through the dataset). To prevent overfitting, we used a cross-validation approach in which 10% of the data were held out (i.e., not used for training) at each iteration and used to evaluate the loss (equation 2). We implemented early stopping in which training stopped if the validation loss did not improve after three iterations. Models were implemented using the Keras API of TensorFlow in Python (Chollet, 2015). As mentioned above, the model produces two sets of species vectors (*W* and 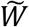), but since the species co-occurrence matrices are symmetric, the two sets are equivalent and differ only due to their random initializations. Following Pennington *et al*. (2014), we took the elementwise averages of the two sets, resulting in a single *m*-dimensional vector *w* and bias term *b* for each species. Overall, we created 20 sets of species vectors (global or local x 5 vector dimensions x 2 weighting function cutoffs). Bias terms are usually not included in word embeddings, but we were interested in testing whether they contain additional information that improves prediction. We therefore compared the predictive performance of species vectors alone or with bias terms included.

We compared the performance of species vectors generated by GloVe with those generated by tb-PCA (Legendre & Gallagher, 2001), an ordination method widely used in ecology. Due to the large size of the sPlotOpen site-by-species matrix (>3.6 billion entries), most traditional ordination methods, including standard PCA, were not computationally feasible. We used a partial PCA method, FlashPCA (Abraham et al., 2017), which performs a partial decomposition with a specified number of principle components and scales to very large datasets. We perform partial PCA on the Hellinger-transformed site-by-species matrix with 4, 8, 16, 32, or 64 principal components.

#### Relationship with functional traits and phylogeny

To assess the degree to which the information encoded in species vectors learned from co-occurrence data overlaps with functional trait and phylogenetic information, we obtained a gap-filled dataset of 18 traits representing leaf, wood, and seed characteristics based on trait records from the TRY database (Kattge et al., 2020) and using the BHPMF gap-filling algorithm (Schrodt et al., 2015) for 21,857 species in sPlotOpen (Bruelheide et al., 2019; Sabatini et al., 2021). We created a phylogeny based on Smith & Brown’s (2018) phylogeny of seed plants using the phylo.maker function in the V.PhyloMaker package in R (Jin & Qian, 2019). We cleaned and harmonized species names following the World Flora Online taxonomic backbone using the WorldFlora package (Kindt, 2020), resulting in a data set of 20,365 species. We calculated pairwise Euclidean distances between species based on species vectors and (log-transformed, scaled) functional traits. We assessed the relationships between species vector, trait, and phylogenetic distances using Mantel tests with 999 permutations.

#### Test of predictive performance

We tested the ability of species vectors learned from co-occurrence data to predict elevational range shifts of plant species in European mountain ranges using the dataset of Lenoir *et al*. (2008). The dataset consists of the optimum elevation (altitude at maximum probability of occurrence) of 171 forest plant species for the periods 1905-1985 and 1986-2005. The elevational range shift was calculated as the difference in optimum elevation (in meters) between the two periods. We aimed to compare the predictive performance of co-occurrence-based species vectors with that of functional traits and phylogenetic information. We matched species across the range shift, co-occurrence, trait, and phylogenetic data, resulting in a data set of 162 species used to model range shifts. To incorporate phylogenetic information, we calculated the pairwise cophenetic phylogenetic distance matrix between species and extracted phylogenetic eigenvectors (Diniz-Filho et al., 1998) using the PVRdecomp function in the PVR package in R (Santos, 2018).

To predict species range shifts, we fit regularized linear models with species vectors, traits, phylogenetic eigenvectors, species vectors plus traits, or species vectors plus phylogenetic eigenvectors as the predictors and range shift (in m) as the response variable. Regularization reduces overfitting, a concern given the relatively small sample size and large number of predictors in the models, by penalizing large regression coefficients. We used the elastic net penalty (Zou & Hastie, 2005), which mixes ridge (penalizes the square of coefficients, tending to shrink them) and lasso (penalizes the absolute value of coefficients, tending to keep some and drop others, effectively performing variable selection) regularization, with the ratio controlled by the parameter α (α = 0, ridge; α = 1, lasso) and overall strength of penalization controlled by parameter λ. To select the optimum values of these parameters, we conducted a grid search using five values of α ranging from 0 to 1 and 10 values of λ ranging from 0.018 to 148. For each parameter combination, we conducted repeated 10-fold cross-validation (10 repeats) and selected the parameter values that had the lowest root mean squared error (RMSE) averaged across the hold-out sets (i.e., data not used in training for each fold and repeat). We used this cross-validation grid search procedure to select optimum parameter values and calculate performance metrics (RMSE, R^2^) for each set of predictors. This cross-validation approach estimates the predictive performance of the models on independent data (i.e., species not used in model fitting), which is the key test of the ability of the models to generalize to new species.

#### Interpreting the vectors

To explore the ecological meaning of the species vectors, we identified the vector dimensions that were most important for explaining species range shifts (i.e., those that had the largest standardized regression coefficients) and tested their correlations with species’ functional traits and aspects of their geographic and environmental distributions. We quantified species distributions as the mean latitude, longitude, elevation, vegetation type, biome, and environmental scores (based on a PCA of 30 climate and soil variables) of sPlotOpen sites in which the species occurred.

## RESULTS

### Relationships between co-occurrence-based vectors, functional traits, and phylogeny

Species vectors learned from co-occurrence data using the GloVe algorithm were significantly but weakly correlated with functional traits (Pearson *r* = 0.01-0.07; *P* = 0.001-0.006) and phylogenetic distances (*r* = 0.04-0.07; *P* = 0.001), indicating that the co-occurrence-based vectors encode information largely not captured by traits or phylogeny. By comparison, traits and phylogenetic distances were much more strongly correlated with each other (*r* = 0.25; *P* = 0.001), indicating a moderately strong phylogenetic signal in traits.

### Predictive performance

Species vectors generated by GloVe performed considerably better than functional traits and phylogenetic eigenvectors at predicting species’ elevational range shifts (Fig. 2). Overall, the best models using species vectors alone or in combination with traits explained 20.2% of variance in range shifts of out-of-sample species, compared to 10.9% variance explained by the best model using traits alone (Fig. 2; Table S1). Species vectors and traits together generally performed similarly to species vectors alone. No variables were selected in the best model using phylogenetic eigenvectors (i.e., all coefficients were shrunk to zero), indicating that phylogenetic information had little to no predictive value. We obtained similar results using models that only considered subsets of leading eigenvectors. Including phylogenetic eigenvectors together with co-occurrence-based species vectors reduced performance relative to models with species vectors alone (Table S1). We confirmed the lack of predictive power of phylogenetic information by directly testing for phylogenetic signal in species’ range shifts, finding that there was essentially none (Pagel’s λ < 0.001, *P* = 1).

We tested how the performance of species vectors generated by GloVe was affected by the training data set (“global” vs. “local”), vector length, weighting threshold, and inclusion of bias terms (Fig. 2; Table S1). These factors affected performance interactively, but vector length had the largest effect, with predictive accuracy generally peaking at intermediate sizes (8-16 dimensions for global vectors, 32 dimensions for local vectors). Global vectors (trained on all plots in the sPlotOpen data set) and local vectors (trained only on plots that contained at least one species in the range shifts data set) had similar performance. Weighting threshold and inclusion of bias terms generally had weak effects on performance (Fig. 2; Table S1).

**Figure 2.**
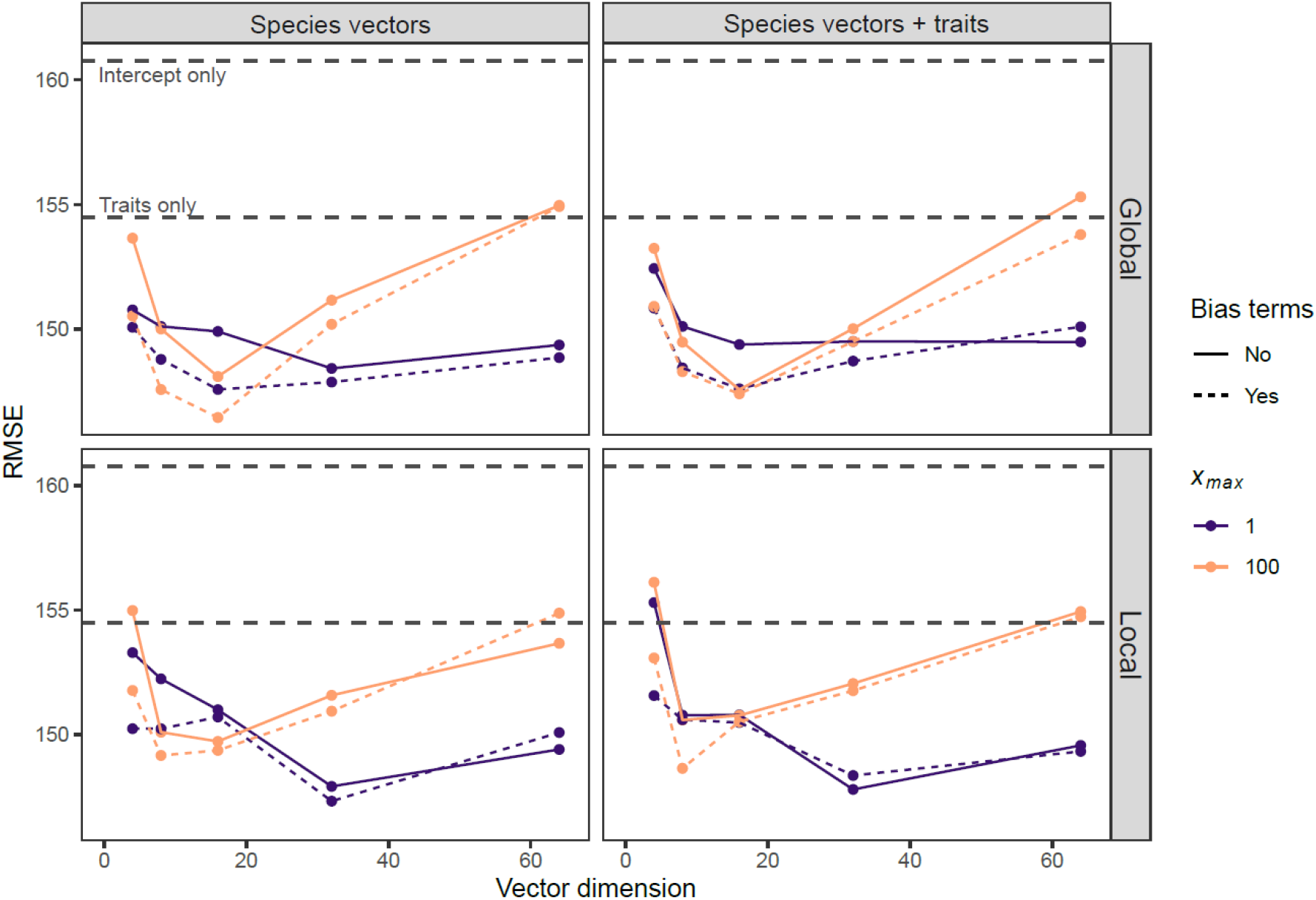
Performance of co-occurrence-based species vectors generated by GloVe and traits at predicting elevational range shifts of 162 European plant species. Predictive performance, measured as the average RMSE on out-of-sample data from repeated 10-fold cross validation, was compared for models including species vectors only (left column) or species vectors plus traits (right column). Species vectors were trained on “global” (all plots in sPlotOpen dataset; top row) or “local” (only plots that contain at least one species in the range shifts dataset; bottom row) data. Performance of species vectors was assessed as a function of vector dimension, inclusion of bias terms, and weighting function threshold (*x_max_*). Horizontal gray lines show the performance of the intercept-only and traits-only models.

Species vectors generated by tb-PCA outperformed traits and phylogenetic eigenvectors at predicting species range shifts but did not explain as much variance as vectors generated by GloVe (Fig. 3; Table S1). Performance of vectors generated by tb-PCA decreased with increasing vector dimension (Fig. 3).

**Figure 3.**
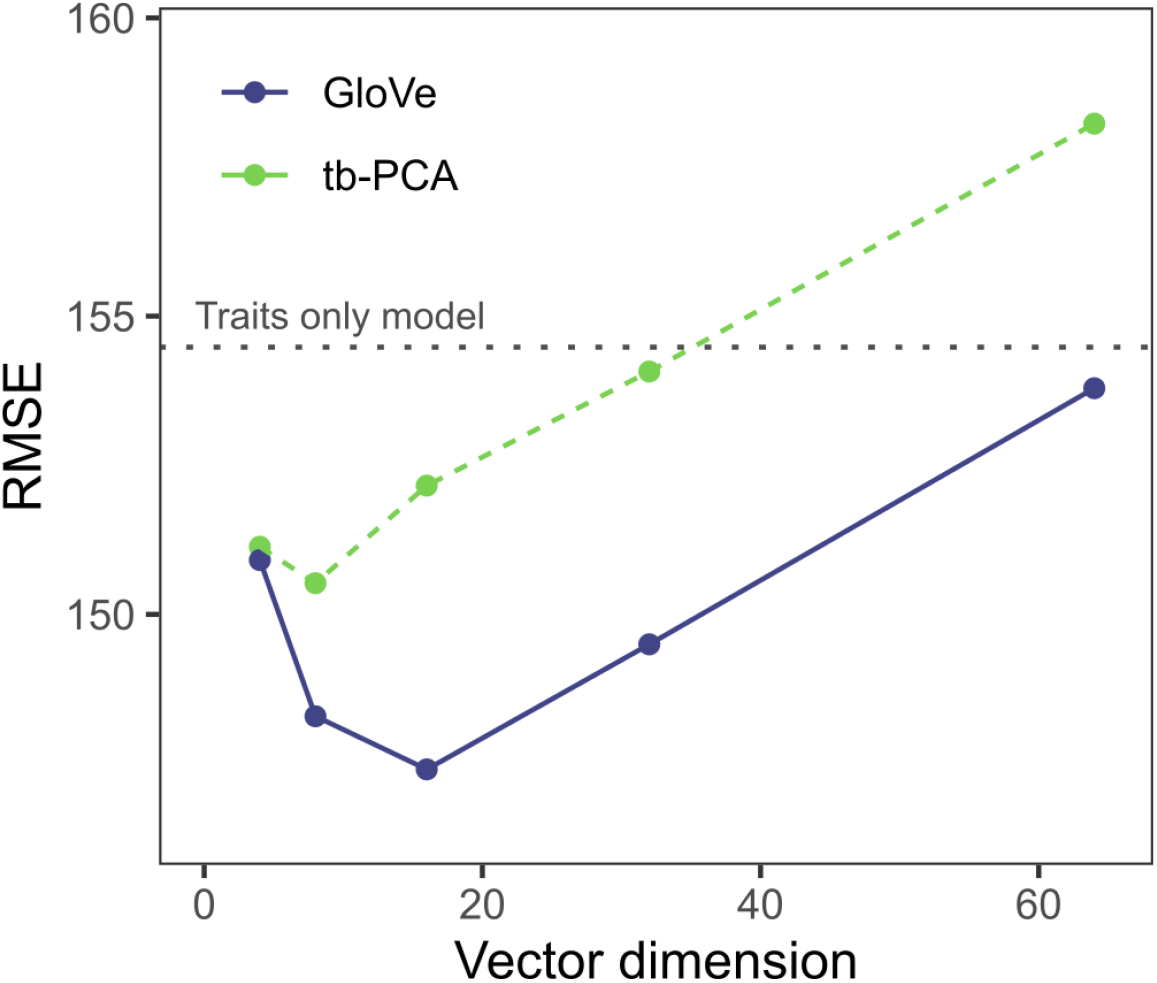
Comparison of performance of species vectors generated by GloVe (vectors trained on global data with bias terms included, *X_max_* = 100) and transformation based-PCA at predicting elevational range shifts of 162 European plant species. Predictive performance was measured as the average root mean squared error (RMSE) of predictions for out-of-sample data from repeated 10-fold cross validation.

#### Ecological interpretation

We explored the ecological correlates of vector dimensions 1 and 8, which were the most important dimensions (i.e., had the largest standardized regression coefficients; Table S2) for explaining range shifts out of the set of vectors that had the best performance in our tests (global vectors with 16 dimensions, bias terms included, *Xmax* = 100; Table S1). Dimension 1 was associated with multiple functional traits (Fig 4; Table S3). Species with high values on dimension 1 tended to have large upward shifts in elevation, fast leaf economics traits (high SLA and leaf P, low LDMC), large leaves, small seeds, and low-density stems. Globally, they tended to be associated with wet, cold, high-elevation, forested habitats (Table S4). Dimension 8 mostly captured aspects of species’ geographic distributions and was weakly correlated with functional traits (Fig. 4; Table S3, S4). Species with low values on dimension 8 tended to have large upward range shifts and were most likely to occur in boreal and dry midlatitude biomes, with distributions centered in eastern Eurasia (high longitudes).

**Figure 4.**
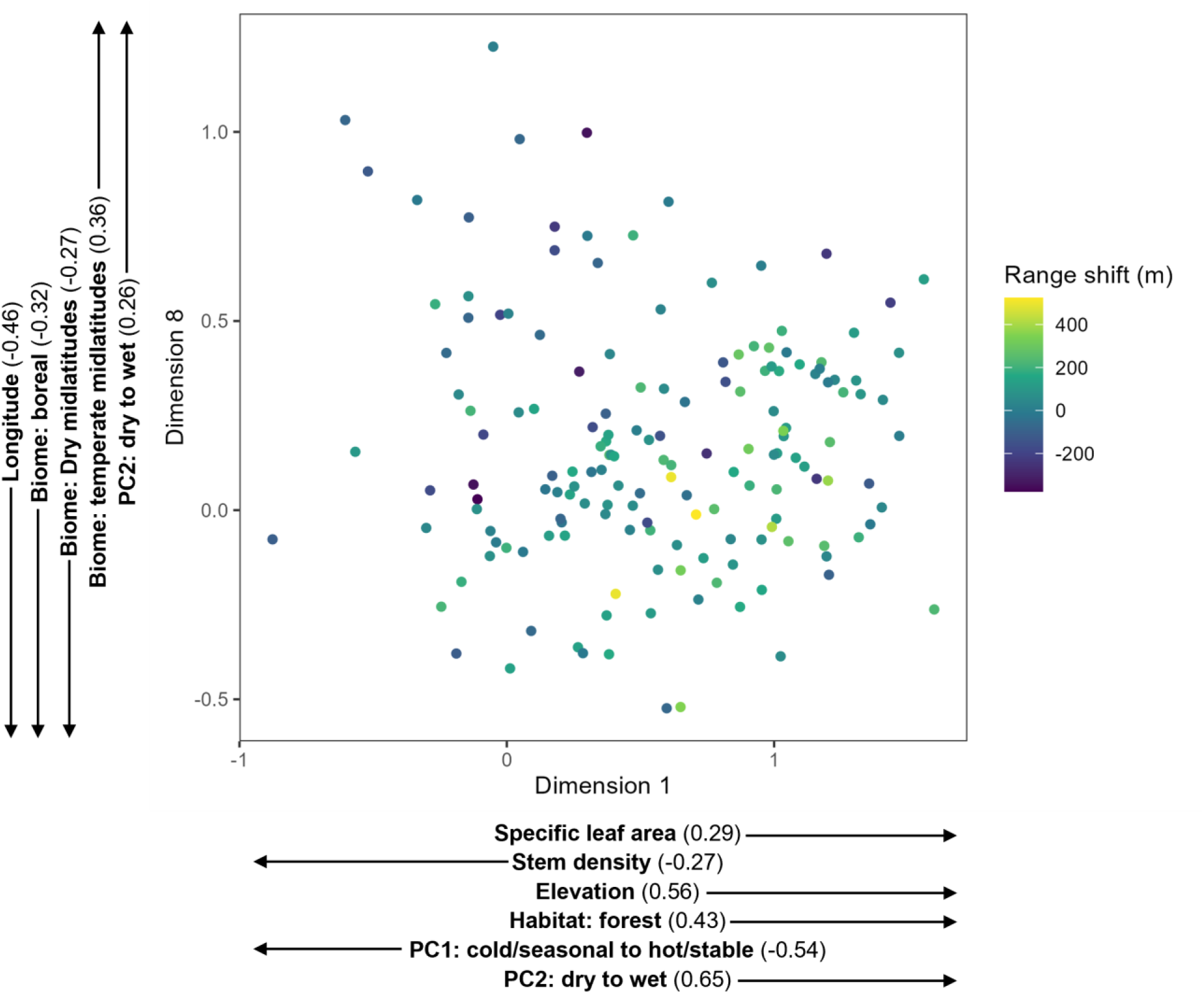
Locations of European montane plant species in co-occurrence-based vector space. Species are plotted in the two dimensions that explained the most variation in species range shifts out of the set of vectors with the best performance in our tests (GloVe vectors trained on global sPlotOpen data with bias terms included, *X_max_* = 100). Color indicates the magnitude of species range shifts. Arrows show the direction of correlation with functional traits and biogeographic attributes that were significantly correlated with each vector dimension (Pearson correlation coefficients are shown in parentheses).

## DISCUSSION

Here we argue that species co-occurrence patterns contain valuable information about species’ ecological attributes, and that harnessing this information could improve ecological prediction. Inspired by methods used in language modeling (Mikolov et al., 2013; Pennington et al., 2014), we present an efficient, scalable method for encoding information contained in co-occurrence patterns in vector representations of species. We demonstrate this method by generating vectors for nearly 40,000 plant species using a global vegetation dataset, and we show that the vectors outperform traits and phylogenetic information in a case study predicting elevational range shifts in European montane plant species. Given the widespread availability of species occurrence data, species vectors learned from co-occurrence patterns are a widely applicable tool for capturing information about the ecological attributes of species, with many potential applications for describing and predicting the ecology of species, communities, and ecosystems.

In theory, any ordination method could be used to derive species vectors from ecological community data, but in practice most existing ordination methods used in ecology cannot handle very large datasets. We compared the performance of two methods, GloVe and tb-PCA, that could scale to the global vegetation dataset in our case study. GloVe outperformed tb-PCA, but the comparison is complicated by the large number of parameters that could influence model performance. Vector dimension in particular had a large effect on performance, which peaked at intermediate vector sizes (8-32 dimensions) for GloVe and small sizes (4-8 dimensions) for tb-PCA. Larger vectors likely contain more ecological information, but increasing the number of predictors in a model increases the risk of overfitting, which may explain why the larger vectors had relatively poor predictive performance. The performance of GloVe in language modeling tasks increases monotonically with the amount of training data (Pennington et al., 2014), and it is reasonable to think that, all else being equal, the same applies for ecological data. We compared the performance of GloVe trained on the full global vegetation dataset and on a “local” subset of data that only included sites containing at least one species in our prediction dataset. The global and local vectors had similar performance, indicating that the benefits of “more” vs. “more focused” data mostly cancelled out in this case. We tested the effects of two additional GloVe tuning parameters (bias term inclusion and weighting function threshold) but found they had minimal effects on predictive performance. Nevertheless, we feel that it is not possible to make general recommendations regarding tuning parameters based on a single case study, and we recommend that future users compare different vector dimensions and hyperparameter combinations using cross-validation to find the values that maximize performance in their specific prediction tasks.

Our case study demonstrates that species vectors learned from co-occurrences have predictive power and therefore encode ecological information, but what kind of information? Since the vectors are trained on species co-occurrence patterns, they potentially encode information about any mechanism that influences those patterns (Fig. 1) and will depend on the specifics of the training data and model. Species representations derived from community data collected over a broad spatial extent encompassing multiple biogeographic regions and strong environmental gradients will likely primarily encode information about mechanisms that drive species distributions at broad scales, such as climatic tolerances, evolutionary history, and dispersal (MacArthur, 1972). In contrast, vectors trained on fine-grained community data will be more likely to encode information about species’ attributes that drive their fine-scale distributions, such as edaphic and microhabitat preferences (Kruckeberg, 2004; Whittaker, 1960) and species interactions (Elton, 1946). While we argue that the strength of co-occurrence-based species vectors lies mainly in their predictive performance rather than interpretability, it may be possible to interpret their ecological meaning by examining relationships between vector dimensions and other measured species attributes—e.g., functional traits, range centroids and limits, environmental tolerances or optima. For example, in our case study, we found that the vector dimension that was most important for predicting species range shifts was strongly associated with leaf economics and size traits, indicating that it encodes information about species’ functional strategies. The second most important dimension was unrelated to traits but was correlated with species’ geographic range centers and biome affiliations, suggesting that it encodes information about biogeographic processes that drive species distributions across broad scales. While interpreting the vectors in light of measured traits and environmental associations is useful for understanding their ecological meaning, we believe that the real power of the vectors is that they encode information about unmeasured traits and attributes.

### Applications

In the case study presented here, we focus on using species vectors to predict species range shifts in response to environmental change, but there are many other potential applications. To the extent that the species vectors encode information about the ecological attributes that shape species’ co-occurrence patterns, they could be used to predict virtually anything about a species. Examples include demographic performance, population dynamics, invasiveness, extinction risk, biotic associations and interaction strengths, and effects on ecosystem processes. The key assumption is that species that have similar co-occurrence patterns—and are therefore close to each other in the vector space—have similar values of the prediction target. This means that species vectors will only be useful for making predictions about a given species if observations of the prediction target (e.g., range shifts) are already available for species that are nearby in (at least some dimensions of) the vector space.

In addition to making predictions about individual species, co-occurrence-based species vectors may be useful for characterizing whole communities. Analogous to functional diversity metrics calculated using species’ trait values (Petchey & Gaston, 2002; Villéger et al., 2008), measures of the spread of species in the vector space could quantify hidden dimensions of diversity within and among communities. For example, if species vectors capture differences in species’ resource use or environmental responses that are not easily quantified by traits, measures of diversity based on species vectors may be more predictive of community productivity or stability than measures based on functional traits. Measures of the diversity and distinctiveness of communities in the vector space could help identify priorities for conservation (Cadotte & Davies, 2010; Faith, 1992). Quantifying the locations of undisturbed reference communities in the vector space may help create targets for ecosystem restoration projects (Engst et al., 2016; White & Walker, 1997). In sum, co-occurrence-based vectors can be used alongside functional traits and evolutionary relatedness to quantify additional dimensions of biodiversity, which can be used in multiple applications, within and beyond the plant realm.

### Relationship to other approaches

Ordination methods are commonly used in ecology to reduce the dimensionality of community data, resulting in representations of species in low-dimensional space. Our approach is novel in that 1) we propose using species ordination scores (which we refer to as species vectors) derived from a single (ideally very large) community dataset to represent species in a variety of external prediction tasks involving independent data, and 2) we apply an ordination algorithm, GloVe, that has not previously been used in ecology. Among the ecological ordination methods we are familiar with, GloVe is most similar to model-based ordination methods, such as generalized linear latent variable models (Hui et al., 2015; Warton et al., 2015), which derive species ordination scores (factor loadings) in the course of predicting species abundances. Similarly, GloVe derives species ordination scores in the course of predicting species co-occurrences. GloVe has the advantage of being able to scale to large datasets that most ordination methods used in ecology are unable to handle, as demonstrated by its use in language processing to generate word embeddings from word co-occurrence matrices with billions of entries. Other authors have also had the idea of adapting word embedding algorithms to ecological data. Angelov (2018) used the word2vec algorithm (Mikolov et al., 2013) to generate species embeddings for mammals using a global species occurrence dataset but did not test the ability of the embeddings to make predictions about species. Chen et al. (2017) and Joseph (2020) developed species distribution models that incorporated species embeddings via deep neural networks but did not propose transferring the embeddings to external prediction tasks.

### Extensions and limitations

There are several possible routes for extending and improving our method. First, and most obvious, is training the models on more data. The number of ecological attributes that influence species distributions—and therefore dimensionality of the vector space required to fully capture those attributes—is likely large, so a large amount of co-occurrence data is probably needed to locate species in that high-dimensional space. We suspect that detecting the signals of species interactions will require especially large data sets and high-dimensional vectors. Second, whereas we trained our models on non-zero entries of species co-occurrence matrices, the zeroes (i.e., species pairs that never co-occur) likely contain additional ecological information, and including them in the training data could improve the predictive performance of the resulting species vectors, though this would come at the cost of drastically increased training time if the species co-occurrence matrix were very sparse. Two species might have zero co-occurrences for different reasons: e.g., non-overlapping geographical ranges, habitat discrimination, or competitive exclusion. Designing an efficient approach for including zeroes in the training data therefore requires a strategy for discriminating these different zeroes. For instance, one might decide to only include zeroes for species pairs that have overlapping geographic ranges. Third, there are multiple hyperparameters of the GloVe algorithm, optimization routine, and predictive model that could be tuned more finely. Finally, different algorithms could be used to learn the species vectors. We chose GloVe due to its record of success in natural language processing and ability to handle huge amounts of data, but other algorithms, including ordination methods traditionally used in ecology, might also prove effective if computational challenges that limit their application to very large datasets can be overcome. Development and comparison of other ordination methods for generating species vectors for ecological prediction should be a priority for future research.

The most obvious limitation of our approach is that the species vectors may be difficult to interpret. This means that even if they are valuable for prediction, they may be less helpful for understanding mechanisms. Another limitation is that the approach is data hungry. The information content of the vectors scales with the amount of training data, and they will likely be most transferrable when the co-occurrence data used to derive the vectors includes species and environments that overlap with the prediction task. Training the vectors on co-occurrence datasets that include observations collected over a large spatial and temporal extent will likely maximize their ecological information content and increase their transferability. If co-occurrence data are scarce, as is likely the case with rare species, the vectors will have limited predictive value. Similarly, species vectors trained on co-occurrence data collected in a particular environment may not transfer well to making predictions in different environmental contexts. Finally, while our approach can be applied to any system for which species occurrence data are available, we suspect it will be most effective for sessile organisms, whose co-occurrence patterns are likely more stable and may contain more ecological information than those of highly mobile organisms.

### Conclusions

If ecologists are going to have any chance of success at making predictions beyond the handful of species we are able to study directly in any given context, we need to base our predictions on the ecological attributes that underly species’ responses to and effects on their environment (Buckley & Kingsolver, 2012; Lavorel & Garnier, 2002). This need has motivated trait-based and phylogenetic approaches in ecology, on the basis that traits and phylogenetic relationships carry useful information about species’ ecology that go beyond individual species contingencies. Here we argue that co-occurrence patterns are another valuable source of information and present a method for harnessing it. Our chance of success at making generalizable predictions across many species will likely be maximized if we take advantage of multiple sources of information to describe species, including traits, phylogeny, and co-occurrence patterns. Our approach also illustrates the potential of transfer learning—taking information learned from one context and applying to others (Pan & Yang, 2010)—to improve ecological predictions. We have made species vectors learned from co-occurrences for nearly 40,000 plant species available (see data availability statement), and we encourage researchers in plant ecology to use them in their prediction tasks. We now have huge amounts of data in ecology and examples from other fields of how to leverage the information in high-dimensional datasets to make predictions (Farley et al., 2018). Unsupervised and transfer learning approaches such as the one we describe in this paper can be an important tool in our toolkit, together with traditional statistical and mechanistic models, for improving prediction in ecology.

## Acknowledgements

We thank Jens Kattge for making the gap-filled trait data available and providing feedback on a draft of the manuscript. This study was funded by the National Science Foundation EPSCOR Grant #2019528. Francesco Maria Sabatini gratefully acknowledges the support of the Italian Ministry of University and Research, under the Maria Levi Montalcini programme (2019).

## Author contributions

AS and DCL conceived the ideas and designed methodology; AS and FMS assembled the data; AS analysed the data; AS led the writing of the manuscript. All authors contributed critically to the drafts and gave final approval for publication.

## Data availability

Data and code to reproduce the results presented in this paper are available at: https://github.com/andrewsiefert/speciesVectors.

## References

Abraham, G., Qiu, Y., & Inouye, M. (2017). FlashPCA2: principal component analysis of Biobank-scale genotype datasets. Bioinformatics, 33, 2776–2778. https://doi.org/10.1093/bioinformatics/btx299

Angelov, B. (2018). species2vec: A novel method for species representation. BioRxiv:461996. https://doi.org/10.1101/461996

Angert, A. L., Crozier, L. G., Rissler, L. J., Gilman, S. E., Tewksbury, J. J., & Chunco, A. J. (2011). Do species traits predict recent shifts at expanding range edges? Ecology Letters, 677–689. https://doi.org/10.1111/j.1461-0248.2011.01620.x

Beissinger, S. R., & Riddell, E. A. (2021). Why are species’ traits weak predictors of range shifts? Annual Review of Ecology, Evolution, and Systematics, 52, 47–66. https://doi.org/10.1146/annurev-ecolsys-012021-092849

Bruelheide, H., Dengler, J., Jiménez-Alfaro, B., Purschke, O., Hennekens, S. M., Chytrý, M., Pillar, V. D., Jansen, F., Kattge, J., Sandel, B., Aubin, I., Biurrun, I., Field, R., Haider, S., Jandt, U., Lenoir, J., Peet, R. K., Peyre, G., Sabatini, F. M., … Zverev, A. (2019). sPlot – A new tool for global vegetation analyses. Journal of Vegetation Science, 30(2), 161–186. https://doi.org/10.1111/jvs.12710

Buckley, L. B., & Kingsolver, J. G. (2012). Functional and phylogenetic approaches to forecasting species’ responses to climate change. Annual Review of Ecology, Evolution, and Systematics, 43(1), 205–226. https://doi.org/10.1146/annurev-ecolsys-110411-160516

Cadotte, M., Albert, C. H., & Walker, S. C. (2013). The ecology of differences: assessing community assembly with trait and evolutionary distances. Ecology Letters, 16(10), 1234–1244. https://doi.org/10.1111/ele.12161

Cadotte, M. W., & Davies, T. J. (2010). Rarest of the rare: Advances in combining evolutionary distinctiveness and scarcity to inform conservation at biogeographical scales. Diversity and Distributions, 16(3), 376–385. https://doi.org/10.1111/j.1472-4642.2010.00650.x

Carmona, C. P., & Pärtel, M. (2021). Estimating probabilistic site-specific species pools and dark diversity from co-occurrence data. Global Ecology and Biogeography, 30(1), 316–326. https://doi.org/10.1111/geb.13203

Chen, D., Xue, Y., Fink, D., Chen, S., & Gomes, C. P. (2017). Deep multi-species embedding. arXiv:1609.09353. https://doi.org/10.24963/ijcai.2017/509

Chollet, F. (2015). Keras. https://keras.io

Collobert, R., Weston, J., Bottou, L., Karlen, M., Kavukcuoglu, K., & Kuksa, P. (2011). Natural language processing (almost) from scratch. Journal of Machine Learning Research, 12, 2493–2537.

Connor, E. F., & Simberloff, D. (1979). The assembly of species communities: chance or competition? Ecology, 60(6), 1132–1140.

Diamond, J. M. (1975). Assembly of species communities. In M. L. Cody & J. M. Diamond (Eds.), Ecology and evolution of communities (pp. 342–444). Harvard University Press.

Diniz-Filho, J. A. F., De Sant’Ana, C. E. R., & Bini, L. M. (1998). An eigenvector method for estimating phylogenetic inertia. Evolution, 52(5), 1247–1262. https://doi.org/10.1111/j.1558-5646.1998.tb02006.x

Elton, C. (1946). Competition and the structure of ecological communities. Journal of Animal Ecology, 15, 54–68.

Engst, K., Baasch, A., Erfmeier, A., Jandt, U., May, K., Schmiede, R., & Bruelheide, H. (2016). Functional community ecology meets restoration ecology: Assessing the restoration success of alluvial floodplain meadows with functional traits. Journal of Applied Ecology, 53(3), 751–764. https://doi.org/10.1111/1365-2664.12623

Faith, D. P. (1992). Conservation evaluation and phylogenetic diversity. Biological Conservation, 61(1), 1–10.

Farley, S. S., Dawson, A., Goring, S. J., & Williams, J. W. (2018). Situating ecology as a bigdata science: Current advances, challenges, and solutions. BioScience, 68(8), 563–576. https://doi.org/10.1093/biosci/biy068

Firth, J. R. (1957). A synopsis of linguistic theory, 1930-1955. In J. R. Firth (Ed.), Studies in Linguistic Analysis (pp. 1–32). Blackwell.

Funk, J. L., Larson, J. E., Ames, G. M., Butterfield, B. J., Cavender-Bares, J., Firn, J., Laughlin, D. C., Sutton-Grier, A. E., Williams, L., & Wright, J. (2017). Revisiting the Holy Grail: Using plant functional traits to understand ecological processes. Biological Reviews, 92(2), 1156–1173. https://doi.org/10.1111/brv.12275

Gotelli, N. (2000). Null model analysis of species co-occurrence patterns. Ecology, 81(9), 2606–2621. http://www.esajournals.org/doi/abs/10.1890/0012-9658(2000)081[2606:NMAOSC]2.0.CO;2

Guisan, A., & Zimmermann, N. E. (2000). Predictive habitat distribution models in ecology. Ecological Modelling, 135, 147–186. https://doi.org/10.1016/S0304-3800(00)00354-9

Hinton, G. E. (1986). Learning distributed representations of concepts. Proceedings of the Eighth Annual Conference of the Cognitive Science Society, 1–12.

Hortal, J., De Bello, F., Diniz-Filho, J. A. F., Lewinsohn, T. M., Lobo, J. M., & Ladle, R. J. (2015). Seven shortfalls that beset large-scale knowledge of biodiversity. Annual Review of Ecology, Evolution, and Systematics, 46, 523–549. https://doi.org/10.1146/annurev-ecolsys-112414-054400

Hui, F. K. C., Taskinen, S., Pledger, S., Foster, S. D., & Warton, D. I. (2015). Model-based approaches to unconstrained ordination. Methods in Ecology and Evolution, 6(4), 399–411. https://doi.org/10.1111/2041-210X.12236

Jin, Y., & Qian, H. (2019). V.PhyloMaker: an R package that can generate very large phylogenies for vascular plants. Ecography, 42(8), 1353–1359. https://doi.org/10.1111/ecog.04434

Jones, K. E., Bielby, J., Cardillo, M., Fritz, S. A., O’Dell, J., Orme, C. D. L., Safi, K., Sechrest, W., Boakes, E. H., Carbone, C., Connolly, C., Cutts, M. J., Foster, J. K., Grenyer, R., Habib, M., Plaster, C. A., Price, S. A., Rigby, E. A., Rist, J., … Purvis, A. (2009). PanTHERIA: a species-level database of life history, ecology, and geography of extant and recently extinct mammals. Ecology, 90(9), 2648–2648. https://doi.org/10.1890/08-1494.1

Joseph, M. B. (2020). Neural hierarchical models of ecological populations. Ecology Letters, 23(4), 734–747. https://doi.org/10.1111/ele.13462

Kattge, J., Bönisch, G., Díaz, S., Lavorel, S., Prentice, I. C., Leadley, P., Tautenhahn, S., Werner, G. D. A., Aakala, T., Abedi, M., Acosta, A. T. R., Adamidis, G. C., Adamson, K., Aiba, M., Albert, C. H., Alcántara, J. M., Alcázar C, C., Aleixo, I., Ali, H., … Wirth, C. (2020). TRY plant trait database – enhanced coverage and open access. Global Change Biology, 26(1), 119–188. https://doi.org/10.1111/gcb.14904

Kindt, R. (2020). WorldFlora: An R package for exact and fuzzy matching of plant names against the World Flora Online taxonomic backbone data. Applications in Plant Sciences, 8(9), 1–5. https://doi.org/10.1002/aps3.11388

Kruckeberg, A. R. (2004). Geology and plant life: the effects of landforms and rock types on plants. University of Washington Press.

Lavorel, S., & Garnier, E. (2002). Predicting changes in community composition and ecosystem functioning from plant traits: revisiting the Holy Grail. Functional Ecology, 16, 545–556.

Legendre, P., & Gallagher, E. (2001). Ecologically meaningful transformations for ordination of species data. Oecologia, 129(2), 271–280. https://doi.org/10.1007/s004420100716

Lenoir, J., Gégout, J. C., Marquet, P. A., De Ruffray, P., & Brisse, H. (2008). A significant upward shift in plant species optimum elevation during the 20th century. Science, 320(5884), 1768–1771. https://doi.org/10.1126/science.1156831

Lewin, R. (1983). Santa Rosalia was a goat. Science, 221(4611), 636–639.

Li, Y., & Yang, T. (2018). Word embedding for understanding natural language: a survey. In S. Srinivasan (Ed.), Guide to Big Data Applications (pp. 83–104). Springer.

MacArthur, R. H. (1972). Geographical ecology: patterns in the distribution of species. Princeton University Press.

Madin, J. S., Anderson, K. D., Andreasen, M. H., Bridge, T. C. L., Cairns, S. D., Connolly, S. R., Darling, E. S., Diaz, M., Falster, D. S., Franklin, E. C., & others. (2016). The Coral Trait Database, a curated database of trait information for coral species from the global oceans. Scientific Data, 3(1), 1–22.

McGill, B. J., Enquist, B. J., Weiher, E., & Westoby, M. (2006). Rebuilding community ecology from functional traits. Trends in Ecology and Evolution, 21(4), 178–185. https://doi.org/10.1016/j.tree.2006.02.002

Mikolov, T., Chen, K., Corrado, G., & Dean, J. (2013). Efficient estimation of word representations in vector space. arXiv:1301.3781.

Paine, C. E. T., Amissah, L., Auge, H., Baraloto, C., Baruffol, M., Bourland, N., Bruelheide, H., Daïnou, K., de Gouvenain, R. C., Doucet, J. L., Doust, S., Fine, P. V. A., Fortunel, C., Haase, J., Holl, K. D., Jactel, H., Li, X., Kitajima, K., Koricheva, J., … Hector, A. (2015). Globally, functional traits are weak predictors of juvenile tree growth, and we do not know why. Journal of Ecology, 103(4), 978–989. https://doi.org/10.1111/1365-2745.12401

Pan, S. J., & Yang, Q. (2010). A survey on transfer learning. IEEE Transactions on Knowledge and Data Engineering, 22(10), 1345–1359.

Pennington, J., Socher, R., & Manning, C. D. (2014). GloVe: Global vectors for word representation. Proceedings of the 2014 Conference on Empirical Methods in Natural Language Processing (EMNLP), 1532–1543.

Peres-Neto, P. R., Olden, J. D., & Jackson, D. A. (2001). Environmentally constrained null models: site suitability as occupancy criterion. Oikos, 93(1), 110–120.

Petchey, O. L., & Gaston, K. J. (2002). Functional diversity (FD), species richness and community composition. Ecology Letters, 5, 402–411.

Sabatini, F. M., Lenoir, J., Hattab, T., Arnst, E. A., Chytrý, M., Dengler, J., De Ruffray, P., Hennekens, S. M., Jandt, U., Jansen, F., Jiménez-Alfaro, B., Kattge, J., Levesley, A., Pillar, V. D., Purschke, O., Sandel, B., Sultana, F., Aavik, T., Aćić, S., … Bates, A. (2021). sPlotOpen – An environmentally balanced, open-access, global dataset of vegetation plots. Global Ecology and Biogeography, December 2020, 1–25. https://doi.org/10.1111/geb.13346

Santos, T. (2018). PVR: Phylogenetic Eigenvectors Regression and Phylogentic Signal-Representation Curve (R package version 0.3). https://cran.r-project.org/package=PVR

Schrodt, F., Kattge, J., Shan, H., Fazayeli, F., Joswig, J., Banerjee, A., Reichstein, M., Bönisch, G., Díaz, S., Dickie, J., Gillison, A., Karpatne, A., Lavorel, S., Leadley, P., Wirth, C. B., Wright, I. J., Wright, S. J., & Reich, P. B. (2015). BHPMF - a hierarchical Bayesian approach to gap-filling and trait prediction for macroecology and functional biogeography. Global Ecology and Biogeography, 24(12), 1510–1521. https://doi.org/10.1111/geb.12335

Smith, S. A., & Brown, J. W. (2018). Constructing a broadly inclusive seed plant phylogeny. American Journal of Botany, 105(3), 302–314. https://doi.org/10.1002/ajb2.1019

Strydom, T., Catchen, M. D., Banville, F., Caron, D., Dansereau, G., Desjardins-Proulx, P., Forero-Muñoz, N. R., Higino, G., Mercier, B., Gonzalez, A., Gravel, D., Pollock, L., & Poisot, T. (2021). A roadmap towards predicting species interaction networks (across space and time). Philosophical Transactions of the Royal Society B: Biological Sciences, 376(1837). https://doi.org/10.1098/rstb.2021.0063

Ulrich, W. (2004). Species co-occurrences and neutral models: reassessing J. M. Diamond’s assembly rules. Oikos, 107, 603–609.

Villéger, S., Mason, N. W. H., & Mouillot, D. (2008). New multidimensional functional diversity indices for a multifaceted framework in functional ecology. Ecology, 89(8), 2290–2301. http://www.esajournals.org/doi/pdf/10.1890/07-1206.1

Warren, D. L., Cardillo, M., Rosauer, D. F., & Bolnick, D. I. (2014). Mistaking geography for biology: Inferring processes from species distributions. Trends in Ecology and Evolution, 29(10), 572–580. https://doi.org/10.1016/j.tree.2014.08.003

Warton, D. I., Blanchet, F. G., O’Hara, R. B., Ovaskainen, O., Taskinen, S., Walker, S. C., & Hui, F. K. C. (2015). So many variables: jointmodeling in community ecology. Trends in Ecology and Evolution, 30(12), 766–779. https://doi.org/10.1016/j.tree.2015.09.007

Webb, C. O., Ackerly, D. D., McPeek, M. A., & Donoghue, M. J. (2002). Phylogenies and community ecology. Annual Review of Ecology and Systematics, 33(1), 475–505. https://doi.org/10.1146/annurev.ecolsys.33.010802.150448

White, P. S., & Walker, J. L. (1997). Approximating nature’s variation: Selecting and using reference information in restoration ecology. Restoration Ecology, 5(4), 338–349. https://doi.org/10.1046/j.1526-100X.1997.00547.x

Whittaker, R. H. (1960). Vegetation of the Siskiyou Mountains, Oregon and California. Ecological Monographs, 30(3), 279–338. https://www.jstor.org/stable/1943563

Zou, H., & Hastie, T. (2005). Regularization and variable selection via the elastic net. Journal of the Royal Statistical Society. Series B: Statistical Methodology, 67(2), 301–320. https://doi.org/10.1111/j.1467-9868.2005.00503.x

