## Supporting Information for "You shall know a species by the company it keeps: leveraging co-occurrence data to improve ecological prediction"

**Table S1.** Tests of ability of co-occurrence-based vectors, traits, and phylogenetic eigenvectors to predict elevational range shifts in 162 European montane plant species. Performance metrics on out-of-sample data are shown for models including intercept only, functional traits (“Traits”), phylogenetic eigenvectors (“Phylo”), species vectors generated by GloVe (“GloVe”), Glove vectors plus traits (“GloVe + Traits”), co-occurrence-based vectors plus phylogenetic eigenvectors (“GloVe + Phylo”), and species vectors generated by transformation-based PCA (“tb-PCA”).

| Predictors | Training data | Vector dimension | $x_{max}$ | Bias terms | RMSE | $R^2$ |
|---|---|---|---|---|---|---|
| Intercept only | - | - | - | - | 160.8 | 0.000 |
| Traits | - | - | - | - | 154.5 | 0.109 |
| Phylo | - | - | - | - | 159.2 | 0.000 |
| GloVe | Global | 4 | 1 | No | 152.4 | 0.182 |
| GloVe | Global | 4 | 1 | Yes | 150.8 | 0.174 |
| GloVe | Global | 4 | 100 | No | 153.2 | 0.148 |
| GloVe | Global | 4 | 100 | Yes | 150.9 | 0.173 |
| GloVe | Global | 8 | 1 | No | 150.1 | 0.178 |
| GloVe | Global | 8 | 1 | Yes | 148.4 | 0.190 |
| GloVe | Global | 8 | 100 | No | 149.5 | 0.177 |
| GloVe | Global | 8 | 100 | Yes | 148.3 | 0.187 |
| GloVe | Global | 16 | 1 | No | 149.4 | 0.193 |
| GloVe | Global | 16 | 1 | Yes | 147.6 | 0.211 |
| GloVe | Global | 16 | 100 | No | 147.6 | 0.202 |
| GloVe | Global | 16 | 100 | Yes | 147.4 | 0.213 |
| GloVe | Global | 32 | 1 | No | 149.5 | 0.178 |
| GloVe | Global | 32 | 1 | Yes | 148.7 | 0.164 |
| GloVe | Global | 32 | 100 | No | 150.0 | 0.171 |
| GloVe | Global | 32 | 100 | Yes | 149.5 | 0.176 |
| GloVe | Global | 64 | 1 | No | 149.5 | 0.182 |
| GloVe | Global | 64 | 1 | Yes | 150.1 | 0.173 |
| GloVe | Global | 64 | 100 | No | 155.3 | 0.125 |
| GloVe | Global | 64 | 100 | Yes | 153.8 | 0.125 |
| GloVe | Local | 4 | 1 | No | 155.3 | 0.124 |
| GloVe | Local | 4 | 1 | Yes | 151.6 | 0.152 |
| GloVe | Local | 4 | 100 | No | 156.1 | 0.090 |
| GloVe | Local | 4 | 100 | Yes | 153.1 | 0.131 |
| GloVe | Local | 8 | 1 | No | 150.8 | 0.163 |
| GloVe | Local | 8 | 1 | Yes | 150.6 | 0.160 |
| GloVe | Local | 8 | 100 | No | 150.6 | 0.164 |
| GloVe | Local | 8 | 100 | Yes | 148.6 | 0.205 |
| GloVe | Local | 16 | 1 | No | 150.8 | 0.160 |
| GloVe | Local | 16 | 1 | Yes | 150.5 | 0.162 |
| GloVe | Local | 16 | 100 | No | 150.8 | 0.152 |
| GloVe | Local | 16 | 100 | Yes | 150.5 | 0.154 |

**Table S1 continued**

| Predictors | Training data | Vector dimension | $x_{max}$ | Bias terms | RMSE | $R^2$ |
|---|---|---|---|---|---|---|
| GloVe | Local | 32 | 1 | No | 147.8 | 0.192 |
| GloVe | Local | 32 | 1 | Yes | 148.4 | 0.186 |
| GloVe | Local | 32 | 100 | No | 152.0 | 0.155 |
| GloVe | Local | 32 | 100 | Yes | 151.8 | 0.143 |
| GloVe | Local | 64 | 1 | No | 149.6 | 0.176 |
| GloVe | Local | 64 | 1 | Yes | 149.3 | 0.169 |
| GloVe | Local | 64 | 100 | No | 154.9 | 0.131 |
| GloVe | Local | 64 | 100 | Yes | 154.7 | 0.136 |
| GloVe + Phylo | Global | 4 | 1 | No | 158.8 | 0.000 |
| GloVe + Phylo | Global | 4 | 1 | Yes | 157.4 | 0.000 |
| GloVe + Phylo | Global | 4 | 100 | No | 159.2 | 0.000 |
| GloVe + Phylo | Global | 4 | 100 | Yes | 157.0 | 0.163 |
| GloVe + Phylo | Global | 8 | 1 | No | 158.3 | 0.000 |
| GloVe + Phylo | Global | 8 | 1 | Yes | 159.2 | 0.000 |
| GloVe + Phylo | Global | 8 | 100 | No | 159.0 | 0.000 |
| GloVe + Phylo | Global | 8 | 100 | Yes | 158.9 | 0.000 |
| GloVe + Phylo | Global | 16 | 1 | No | 158.5 | 0.000 |
| GloVe + Phylo | Global | 16 | 1 | Yes | 158.7 | 0.000 |
| GloVe + Phylo | Global | 16 | 100 | No | 158.9 | 0.000 |
| GloVe + Phylo | Global | 16 | 100 | Yes | 158.7 | 0.000 |
| GloVe + Phylo | Global | 32 | 1 | No | 159.1 | 0.000 |
| GloVe + Phylo | Global | 32 | 1 | Yes | 157.3 | 0.125 |
| GloVe + Phylo | Global | 32 | 100 | No | 159.0 | 0.000 |
| GloVe + Phylo | Global | 32 | 100 | Yes | 159.0 | 0.000 |
| GloVe + Phylo | Global | 64 | 1 | No | 158.4 | 0.000 |
| GloVe + Phylo | Global | 64 | 1 | Yes | 155.0 | 0.179 |
| GloVe + Phylo | Global | 64 | 100 | No | 158.8 | 0.000 |
| GloVe + Phylo | Global | 64 | 100 | Yes | 158.6 | 0.000 |
| GloVe + Phylo | Local | 4 | 1 | No | 159.5 | 0.000 |
| GloVe + Phylo | Local | 4 | 1 | Yes | 156.9 | 0.142 |
| GloVe + Phylo | Local | 4 | 100 | No | 158.8 | 0.000 |
| GloVe + Phylo | Local | 4 | 100 | Yes | 158.9 | 0.000 |
| GloVe + Phylo | Local | 8 | 1 | No | 159.0 | 0.000 |
| GloVe + Phylo | Local | 8 | 1 | Yes | 159.2 | 0.000 |
| GloVe + Phylo | Local | 8 | 100 | No | 158.6 | 0.000 |
| GloVe + Phylo | Local | 8 | 100 | Yes | 158.7 | 0.000 |
| GloVe + Phylo | Local | 16 | 1 | No | 158.2 | 0.000 |
| GloVe + Phylo | Local | 16 | 1 | Yes | 158.8 | 0.000 |
| GloVe + Phylo | Local | 16 | 100 | No | 159.0 | 0.000 |
| GloVe + Phylo | Local | 16 | 100 | Yes | 159.0 | 0.000 |

**Table S1 continued**

| Predictors | Training data | Vector dimension | $x_{max}$ | Bias terms | RMSE | $R^2$ |
|---|---|---|---|---|---|---|
| GloVe + Phylo | Local | 32 | 1 | No | 157.1 | 0.110 |
| GloVe + Phylo | Local | 32 | 1 | Yes | 156.7 | 0.121 |
| GloVe + Phylo | Local | 32 | 100 | No | 158.6 | 0.000 |
| GloVe + Phylo | Local | 32 | 100 | Yes | 158.7 | 0.000 |
| GloVe + Phylo | Local | 64 | 1 | No | 159.1 | 0.000 |
| GloVe + Phylo | Local | 64 | 1 | Yes | 156.9 | 0.099 |
| GloVe + Phylo | Local | 64 | 100 | No | 159.2 | 0.000 |
| GloVe + Phylo | Local | 64 | 100 | Yes | 158.4 | 0.104 |
| GloVe + Traits | Global | 4 | 1 | No | 150.8 | 0.147 |
| GloVe + Traits | Global | 4 | 1 | Yes | 150.1 | 0.167 |
| GloVe + Traits | Global | 4 | 100 | No | 153.6 | 0.114 |
| GloVe + Traits | Global | 4 | 100 | Yes | 150.5 | 0.160 |
| GloVe + Traits | Global | 8 | 1 | No | 150.1 | 0.170 |
| GloVe + Traits | Global | 8 | 1 | Yes | 148.8 | 0.173 |
| GloVe + Traits | Global | 8 | 100 | No | 150.0 | 0.162 |
| GloVe + Traits | Global | 8 | 100 | Yes | 147.6 | 0.187 |
| GloVe + Traits | Global | 16 | 1 | No | 149.9 | 0.179 |
| GloVe + Traits | Global | 16 | 1 | Yes | 147.6 | 0.197 |
| GloVe + Traits | Global | 16 | 100 | No | 148.1 | 0.191 |
| GloVe + Traits | Global | 16 | 100 | Yes | 146.5 | 0.202 |
| GloVe + Traits | Global | 32 | 1 | No | 148.4 | 0.181 |
| GloVe + Traits | Global | 32 | 1 | Yes | 147.9 | 0.177 |
| GloVe + Traits | Global | 32 | 100 | No | 151.2 | 0.151 |
| GloVe + Traits | Global | 32 | 100 | Yes | 150.2 | 0.160 |
| GloVe + Traits | Global | 64 | 1 | No | 149.4 | 0.172 |
| GloVe + Traits | Global | 64 | 1 | Yes | 148.9 | 0.165 |
| GloVe + Traits | Global | 64 | 100 | No | 155.0 | 0.122 |
| GloVe + Traits | Global | 64 | 100 | Yes | 154.9 | 0.126 |
| GloVe + Traits | Local | 4 | 1 | No | 153.3 | 0.117 |
| GloVe + Traits | Local | 4 | 1 | Yes | 150.2 | 0.163 |
| GloVe + Traits | Local | 4 | 100 | No | 155.0 | 0.106 |
| GloVe + Traits | Local | 4 | 100 | Yes | 151.8 | 0.163 |
| GloVe + Traits | Local | 8 | 1 | No | 152.2 | 0.138 |
| GloVe + Traits | Local | 8 | 1 | Yes | 150.2 | 0.157 |
| GloVe + Traits | Local | 8 | 100 | No | 150.1 | 0.151 |
| GloVe + Traits | Local | 8 | 100 | Yes | 149.2 | 0.165 |
| GloVe + Traits | Local | 16 | 1 | No | 151.0 | 0.143 |
| GloVe + Traits | Local | 16 | 1 | Yes | 150.7 | 0.146 |
| GloVe + Traits | Local | 16 | 100 | No | 149.7 | 0.175 |
| GloVe + Traits | Local | 16 | 100 | Yes | 149.4 | 0.168 |

**Table S1 continued**

| Predictors | Training data | Vector dimension | $x_{max}$ | Bias terms | RMSE | $R^2$ |
|---|---|---|---|---|---|---|
| GloVe + Traits | Local | 32 | 1 | No | 147.9 | 0.180 |
| GloVe + Traits | Local | 32 | 1 | Yes | 147.3 | 0.185 |
| GloVe + Traits | Local | 32 | 100 | No | 151.6 | 0.140 |
| GloVe + Traits | Local | 32 | 100 | Yes | 150.9 | 0.158 |
| GloVe + Traits | Local | 64 | 1 | No | 149.4 | 0.165 |
| GloVe + Traits | Local | 64 | 1 | Yes | 150.1 | 0.155 |
| GloVe + Traits | Local | 64 | 100 | No | 153.7 | 0.128 |
| GloVe + Traits | Local | 64 | 100 | Yes | 154.9 | 0.117 |
| tb-PCA | Global | 4 | - | - | 151.1 | 0.147 |
| tb-PCA | Global | 8 | - | - | 150.5 | 0.146 |
| tb-PCA | Global | 16 | - | - | 152.2 | 0.142 |
| tb-PCA | Global | 32 | - | - | 154.1 | 0.121 |
| tb-PCA | Global | 64 | - | - | 158.2 | 0.075 |

**Table S2.** Standardized regression coefficients from regularized (elastic net) linear regression of plant species range shifts on GloVe species vectors (global vectors with 16 dimensions, bias terms included, $x_{max}$ = 100). The regression model was fit to the full dataset of European montane plant species (n = 162) using elastic net penalty parameters ($\alpha$ = 0.5, $\lambda$ = 20.1) that maximized out-of-sample predictive accuracy as estimated by repeated 10-fold cross-validation.

| Parameter | Estimate |
|---|---:|
| Intercept | 0.40 |
| Dimension 1 | 0.14 |
| Dimension 2 | 0.00 |
| Dimension 3 | 0.00 |
| Dimension 4 | 0.00 |
| Dimension 5 | 0.00 |
| Dimension 6 | 0.01 |
| Dimension 7 | 0.00 |
| Dimension 8 | -0.17 |
| Dimension 9 | -0.06 |
| Dimension 10 | 0.00 |
| Dimension 11 | -0.01 |
| Dimension 12 | 0.00 |
| Dimension 13 | 0.00 |
| Dimension 14 | 0.00 |
| Dimension 15 | 0.00 |
| Dimension 16 | -0.03 |
| Bias | -0.17 |

**Table S3.** Pearson correlation coefficients for relationships between species vectors generated by GloVe (global vectors with 16 dimensions, bias terms included, $x_{max}$ = 100) and functional traits for species in the European mountains range shifts dataset (n = 162). Results are shown for the two vector dimensions that were most important (largest standardized regression coefficients) for predicting species range shifts. Statistically significant relationships ($P$ < 0.05) are indicated by bold type.

| Variable | Dimension 1 | Dimension 8 |
|---|---|---|
| Leaf area | **0.31** | 0.04 |
| Stem density | **-0.27** | 0.01 |
| Specific leaf area | **0.29** | -0.08 |
| Leaf C per dry mass | -0.10 | -0.08 |
| Leaf N per mass | -0.03 | -0.09 |
| Leaf P per mass | **0.26** | 0.02 |
| Plant height | -0.12 | 0.07 |
| Seed mass | **-0.18** | -0.02 |
| Seed length | -0.03 | -0.02 |
| Leaf dry matter content | **-0.25** | 0.01 |
| Leaf N per area | **-0.36** | 0.04 |
| Leaf N:P | **-0.20** | -0.07 |
| Leaf δ15N | 0.14 | 0.02 |
| Seed number per reproductive unit | **0.20** | 0.08 |
| Leaf fresh mass | **0.23** | 0.06 |
| Stem conduit density | 0.07 | **0.16** |
| Dispersal unit length | -0.03 | -0.08 |
| Wood vessel length | 0.01 | -0.01 |

**Table S4.** Pearson correlation coefficients for relationships between species vectors generated by GloVe (global vectors with 16 dimensions, bias terms included, $x_{max}$ = 100) and biogeographic variables for species in the European mountains range shifts dataset (n = 162). Results are shown for the two vector dimensions that were most important (largest standardized regression coefficients) for predicting species range shifts. Statistically significant relationships ($P$ < 0.05) are indicated by bold type.

| Variable | Dimension 1 | Dimension 8 |
|---|---|---|
| Latitude | **0.16** | 0.01 |
| Longitude | 0.10 | **-0.46** |
| Elevation | **0.56** | -0.06 |
| Habitat: Forest | **0.43** | -0.03 |
| Habitat: Shrubland | **-0.29** | **0.23** |
| Habitat: Grassland | **-0.42** | 0.01 |
| Habitat: Wetland | **-0.33** | 0.13 |
| Biome: Boreal | 0.03 | **-0.32** |
| Biome: Dry midlatitudes | **-0.16** | **-0.27** |
| Biome: Dry tropics and subtropics | 0.06 | **0.19** |
| Biome: Polar and subpolar | -0.07 | -0.08 |
| Biome: Subtropics with winter rain | **-0.41** | **-0.23** |
| Biome: Subtropics with year-round rain | -0.11 | **0.18** |
| Biome: Temperate midlatitudes | **0.25** | **0.36** |
| Biome: Tropics with summer rain | -0.10 | 0.10 |
| Biome: Tropics with year-round rain | 0.05 | -0.05 |
| PC1 (cold/seasonal to hot/stable) | **-0.54** | **0.23** |
| PC2 (dry to wet) | **0.65** | **0.26** |

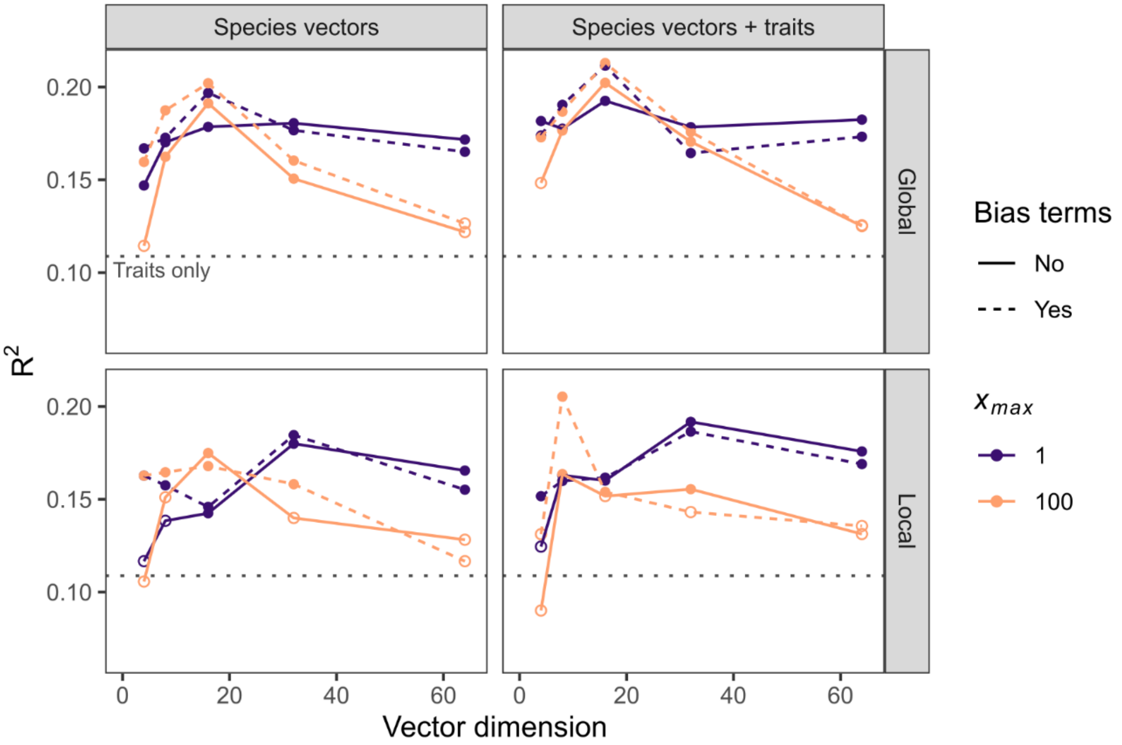


**Figure S1.** Performance of co-occurrence-based species vectors generated by GloVe and traits at predicting elevational range shifts of 162 European plant species. Predictive performance, measured as the average $R^2$ on out-of-sample data from repeated 10-fold cross validation, was compared for models including species vectors only (left column) or species vectors plus traits (right column). Species vectors were trained on "global" (all plots in sPlotOpen dataset; top row) or "local" (only plots that contain at least one species in the range shifts dataset; bottom row) data. Performance of species vectors was assessed as a function of vector dimension, inclusion of bias terms, and weighting function threshold ($x_{max}$). Horizontal dashed line shows the performance of the traits-only model.

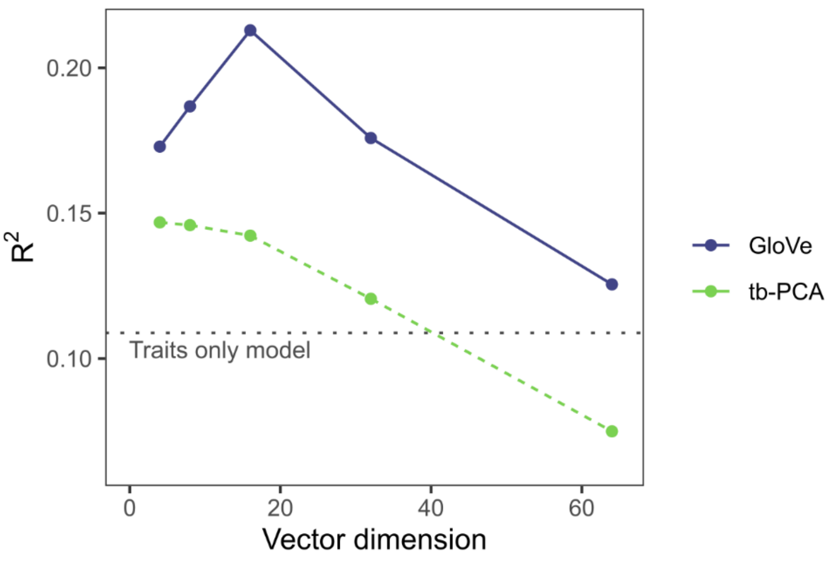


**Figure S2.** Comparison of performance of species vectors generated by GloVe (vectors trained on global data with bias terms included, $x_{max}$ = 100) and transformation based-PCA at predicting elevational range shifts of 162 European plant species. Predictive performance was measured as the average root mean squared error (RMSE) of predictions for out-of-sample data from repeated 10-fold cross validation.
